# Validating a computational framework for ionic electrodiffusion with cortical spreading depression as a case study

**DOI:** 10.1101/2021.11.29.470301

**Authors:** Ada Johanne Ellingsrud, Rune Enger, Didrik Bakke Dukefoss, Geir Halnes, Klas Henning Pettersen, Marie E Rognes

## Abstract

Cortical spreading depression (CSD) is a wave of pronounced depolarization of brain tissue accompanied by substantial shifts in ionic concentrations and cellular swelling. Here, we validate a computational framework for modelling electrical potentials, ionic movement, and cellular swelling in brain tissue during CSD. We consider different model variations representing wild type or knock-out/knock-down mice and systematically compare the numerical results with reports from a selection of experimental studies. We find that the data for several CSD hallmarks obtained computationally, including wave propagation speed, direct current shift duration, peak in extracellular K^+^ concentration as well as a pronounced shrinkage of extracellular space, are well in line with what has previously been observed experimentally. Further, we assess how key model parameters including cellular diffusivity, structural ratios, membrane water and/or K^+^ permeabilities affect the set of CSD characteristics.

**Significance Statement:** Movement of ions and molecules in and between cellular compartments is fundamental for brain function. Cortical spreading depression (CSD) is associated with dramatic failure of brain ion homeostasis. Better understanding the sequence of events in CSD could thus provide new insight into physiological processes in the brain. Despite extensive experimental research over the last decades, even basic questions related to mechanisms underlying CSD remain unanswered. Computational modelling can play an important role going forward, since simulation studies can address hypotheses that are difficult to target experimentally. Here, we assess the physiological validity of a novel mathematical framework for detailed modelling of brain electrodiffusion and osmosis – and provide a platform for in silico studies of CSD and other cerebral electro-mechanical phenomena.

## 1 Introduction

Cortical spreading depression (CSD) is a slowly propagating wave of depolarization of brain cells followed by temporary silencing of electrical brain activity due to a complete collapse of cellular ion homeostasis (Pietrobon & Moskowitz 2014). CSD is characterized by elevated levels of extracellular K^+^ and glutamate (Somjen 2001), swelling of neuronal somata and dendrites (Takano et al. 2007), swelling of astrocyte endfeet (Rosic et al. 2019) and pronounced shrinkage of the extracellular space (ECS) (Mazel et al. 2002). Analyzing the sequence of events in CSD may provide new insight into physiological processes underlying both normal brain function and pathophysiological processes pertinent to a range of brain disorders (Enger et al. 2017).

Despite extensive research over the last decades, even basic questions relating to the mechanisms underlying CSD remain unanswered (Miura et al. 2007). Computational, or *in silico*, modelling may play an important role going forward, not least since simulation studies can address hypotheses and points of debate that are difficult to isolate or address experimentally. Here, we consider a comprehensive computational framework (*the Mori framework*), describing spatial and temporal dynamics of intra- and extracellular ion concentrations, electric potentials, and volume fractions (Mori 2015). The framework has previously been applied to study the roles of glial cells, NMDA receptors and glutamate propagation in CSD (O’Connell & Mori 2016, Tuttle et al. 2019). However, as simulations have not to any significant extent been compared with experimental findings, the physiological validity of the computational framework remains an open question.

To systematically address this issue, we here simulate CSD in different model scenarios, and compare the computational predictions with values from the experimental literature. Our scenarios mimic different mouse models with varying structural and functional parameters, including varying intracellular diffusion, varying transmembrane water and K^+^ permeabilities, and varying membrane characteristics. These choices of model scenarios are in part motivated by the incomplete or disparate findings on the role of AQP4 (Enger et al. 2017, Rosic et al. 2019, Thrane et al. 2013, Yao et al. 2015), and K_ir_4.1 channels (Djukic et al. 2007, Enger et al. 2015, Ohno 2018) in CSD.

Overall, we find that the range of wave speeds, DC shift durations, peak in extracellular K^+^, neuronal changes in volume fraction, and ECS shrinkage obtained computationally all overlap with the experimentally reported ranges in wild type mice. Further, the intracellular glial diffusivity strongly influences the DC shift, while the ratio of neuronal and glial membrane area-to-tissue volume strongly affects the CSD wave speed. Reducing the glial water permeability has a pronounced effect on cellular swelling, whereas the CSD depolarization wavefront speed and the other quantities of interest remain unaltered. In addition, we find that reducing the K_ir_4.1 expression results in reduced glial swelling and depolarization of the glial membrane during CSD.

## 2 Materials and methods

### 2.1 Mathematical and computational framework

We model ionic electrodiffusion and osmotic water flow in brain tissue via the Mori framework; as introduced by Mori (Mori 2015), studied numerically by Ellingsrud et al. (2021), and applied by O’Connell & Mori (2016) and Tuttle et al. (2019). This framework describes tissue dynamics in an arbitrary number of cellular compartments and the ECS via coupled ODEs and PDEs in a model domain. Here, we consider three compartments representing neurons, glia cells and the ECS (see Figure 1), and induce CSD computationally in a one-dimensional model domain of length 10 mm. In particular, we consider the glial subtype astrocyte (for convenience we will henceforth use the terms glia and astrocyte interchangeably). The model predicts the evolution in time and distribution in space of the volume fraction *α*, the electrical potential *ϕ*, and the concentrations [Na^+^], [K^+^], [Cl^−^], [Glu] in each of these compartments. The neuronal and glial membrane potentials are defined as the difference between the neuronal and glial, and extracellular potential, respectively.

**Figure 1.**
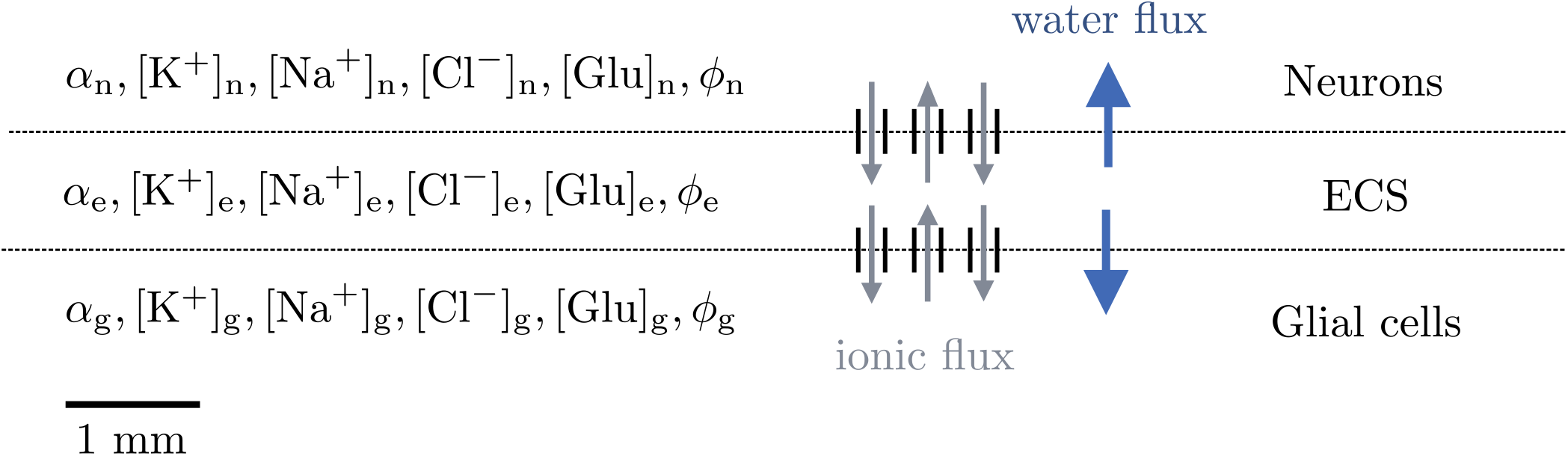
Model overview. The tissue is represented as a 1D domain of length 10 mm including neurons, ECS, and glial cells. Within each compartment, the model describes the dynamics of the volume fraction (*α*), the Na^+^, K^+^, Cl^−^ and glutamate concentrations ([Na^+^], [K^+^], [Cl^−^], [Glu]), and the potential (*ϕ*). Communication between the compartments occur via ionic and/or water membrane fluxes.

Interaction between the compartments is modelled by exchange across the neuron-extracellular and glia-extracellular membranes. In terms of osmotic water flow, the water permeability and area-to-volume ratio of the cellular membranes are key model parameters. To account for transmembrane ion movement across the neuron-extracellular membrane, we consider leak channels, voltage-gated K^+^ and Na^+^ channels, NMDA receptors, and the Na^+^/K^+^-ATPase. Across the glia-extracellular interface, we model leak channels, the inward rectifying K^+^ channel K_ir_4.1, the Na/K/Cl co-transporter, and the Na^+^/K^+^-ATPase. The precise mathematical formulation is detailed in Supplementary Section A, with default parameter values given in Supplementary Section A.3.

### 2.2 Triggers of cortical spreading depression and spreading depolarization

CSD can be triggered in various ways experimentally: by electrical stimulation on the surface of the cerebral cortex (Leao 1944), pinprick (Richter et al. 2002, Rosic et al. 2019), or by topical application of K^+^ (Enger et al. 2017, Yao et al. 2015). SD may occur from conditions associated with ATP depletion and a consequent failure of the Na^+^/K^+^-ATPase, such as ischemia and hypoxia (Cozzolino et al. 2018). All these scenarios results in a local extracellular K^+^ increase, which in turn causes opening of voltage gated cation channels. Here, we consider three different mechanisms for triggering CSD and SD. Specifically, the first two mechanisms trigger CSD, the latter SD:

- Excitatory fluxes: a flux of Na^+^, K^+^ and Cl^−^ is introduced over the neuronal membrane at the first (left-most) 0.02 mm of the computational domain for the first 2 seconds of simulation time.
- Topical application of K^+^: the initial values for the extracellular K^+^ and Cl^−^ concentrations are increased in the first 1 mm of the computational domain.
- Disabled Na^+^/K^+^-ATPase: Na^+^/K^+^-ATPase is disabled by setting the neuron and glia maximum pump rates to zero in the first 1 mm of the computational domain for the first 2 seconds of simulation time.

The precise expressions are detailed in Supplementary Section A.6.

### 2.3 Quantities of interest

Experimental studies of CSD have reported on the speed of CSD waves, the duration and amplitude of extracellular K^+^ and glutamate rises, the DC shift, neuronal and glial swelling, as well as extracellular shrinkage. To compare computational results to the experimental findings, we define the following quantities of interest.

- Mean wave propagation speed (mm/min): given the point *x*_*i*_ at which the neuron potential peaks at time *t*_*i*_, we define the wave speed *v*_*i*_ as *v*_*i*_ = (*x*_*i*_ −*x*_*i*−1_)*/*(*t*_*i*_ −*t*_*i*−1_) at all times *t*_*i*_ for which the neuron potential *ϕ*_n_(*x*_*i*_) has passed a depolarization threshold of −20 mV and after the wave is fully initiated. We then set the mean wave propagation speed as the average of the *v*_*i*_.
- Duration and amplitude of the DC shift: we define the DC shift in terms of the change in extracellular potential as follows. The amplitude of the DC shift is the maximal spatial difference in the extracellular potential (sampled at *t* = 60s). The duration of the DC shift is the difference between the latest and earliest time for which the extracellular potential is below a threshold of −0.05 mV from baseline (sampled at *x* = 1 mm).
- Duration and amplitude of the neuronal, glial and extracellular swelling: these are defined analogously as the duration and amplitude of the DC shift. The lower threshold for neuronal and glial swelling and extracellular shrinkage is set at 0.5%. Shrinkage is defined as negative swelling.
- Duration and amplitude of elevated extracellular K^+^, Cl^−^ and glutamate: these are defined similarly as for the DC shift, with lower thresholds of 8 mM for K^+^ and 002 mM for glutamate, and an upper threshold of 111 mM for Cl^−^.
- Duration and amplitude of neuronal and glial membrane depolarization: these are defined similarly as the above with a lower threshold of −66 mV for the neuronal membrane potential and −77 mV for the glial membrane potential.

In addition to these specific quantities of interest, we plot snapshots in time and the time evolution of the computed fields evaluated at *x* = 1 mm.

### 2.4 Computational model and model variations

As a baseline, we define a wildtype model (Model A, see Table 1) with default model parameters (Table 5 and 6). In addition, we consider four model variations as follows (see also Table 1).

**Table 1.** Overview of the computational models with parameter values: *χ*_g_: glial gap junction factor, *γ*_ne_: neuronal membrane area-to-volume, *γ*_ge_: glial membrane area-to-volume, *η*_ge_: glial membrane water permeability,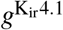: glial K_ir_4.1 resting conductance. Model A corresponds to the default parameters, given in Table 5 and 6 (only 3 significant digits included here). The dash (–) indicates no change from the default values.

| Model | $\chi_g$ | $\gamma_{ne}$ (m<sup>-1</sup>) | $\gamma_{ge}$ (m<sup>-1</sup>) | $\eta_{ge}$ (m<sup>4</sup>/(mol s)) | $g^{K_{ir4.1}}$ (S/m<sup>2</sup>) |
| --- | --- | --- | --- | --- | --- |
| A | 0.05 | $5.38 \times 10^5$ | $6.38 \times 10^5$ | $5.40 \times 10^{-10}$ | 1.30 |
| B | 0 | — | — | — | — |
| C | — | $1.85 \times 10^5$ | $2.00 \times 10^5$ | — | — |
| D | — | — | — | 0 | — |
| E | — | — | — | — | 0.390 |

#### Blocked glial gap junctions

Interconnected astrocytes form syncytia (networks) by gap junctions. Intercellular transport through the astrocytic networks likely facilitate the removal of excess extracellular K^+^ through spatial buffering in the hippocampus (Wallraff et al. 2006). In our computational model, the *glial gap junction factor χ*_*g*_ represents glial gap junctions and defines the effective intercellular diffusion through the astrocytic networks. To explore how diffusion through astrocytic networks affects the CSD wave, we consider a version of the wildtype model with no glial gap junctions (*χ*_g_ = 0) and as a consequence, a zero effective diffusion coefficient in the glial compartment.

#### Reduced membrane area-to-volume

The ratio of membrane area to tissue volume (membrane area-to-volume) for the neuronal and glial compartments are averaged model parameters (*γ*_ne_, *γ*_ge_) that may be difficult to determine experimentally and that are associated with substantial uncertainty. Kager et al. (2000) report an estimated neuronal membrane area-to-volume based on measurements of a rat hippocampal CA1 neuron, and O’Connell & Mori (2016) assume the same value for the glial membrane area-to-volume. Halnes et al. (2019) independently estimate a higher neuronal and glial area-to-volume value. To explore the effect of reducing the neuronal and glial membrane area-to-volume, we consider a model version with reduced area-to-volume parameters (Model C).

#### Reduced glial membrane water permeability

To study the role and effect of water movement across the astrocytic membrane on CSD dynamics, we define a model variant (Model D) by setting the water permeability of the glial membrane *η*_ge_ to 0. This model thus stipulates that water cannot cross the glial membrane; neither via the lipid bilayer itself nor other membrane mechanisms such as AQP4 channels, VRAC, NKCC or other co-transporters. As such, it can be viewed as an extreme case providing an upper bound on the effect of e.g. reduced AQP4 expression on CSD wave characteristics, and allows for comparing computational predictions with the relatively large body of literature on AQP4 in CSD, see e.g. Enger et al. (2017), Rosic et al. (2019), Thrane et al. (2013), Yao et al. (2015). Note that the water permeability is reduced while keeping all other model parameters constant, and as such, model D does account for any potential physiological compensatory mechanisms.

#### *Reduced* K_ir_4.1 *expression*

To study the effect of potassium movement across the astrocytic membrane on the CSD dynamics, we define a model variant (Model E) by reducing the K_ir_4.1 resting conductance of the glial membrane 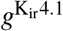. Specifically, we reduce 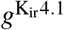 by 70%, and thus represent a partial K_ir_4.1 knockdown. Changes in the membrane parameters affect the steady state of the models, and in response, we also consider a new initial state of the system, see Supplementary Section A.5.

### 2.5 Numerical solution and verification

We apply the solution algorithm presented by (Ellingsrud et al. 2021) and approximate the mathematical model numerically by a finite element method in space, a BDF2 scheme in time and a Strang splitting method. We use spatial and temporal discretization sizes of Δ*x* = 1.25*µ*m and Δ*x* = 3.125 ms, respectively. Numerical verification tests have been carried out to ensure convergence of solutions. The numerical error in the calculated mean wave propagation speed is estimated to be less than 1.5%, whereas for the other quantities of interest we expect the numerical errors to be negligible (Ellingsrud et al. 2021).

### 2.6 Validation principles

We perform a quantitative and qualitative comparison of experimental and computational results via (a selection of the) quantities of interest listed in Section 2.3. We collected experimental findings from a set of recent experimental studies on CSD (Chang et al. 2010, Enger et al. 2017, 2015, Kucharz et al. 2017, Lauritzen & Hansen 1992, Padmawar et al. 2005, Takano et al. 2007, Theis et al. 2003, Thrane et al. 2013, Yao et al. 2015, Zhou et al. 2010). For the comparison, we define intervals of computational and experimental values for each of the quantities of interest reported in (a subset of) the experimental studies (wave speed, DC shift and duration, peak extracellular K^+^ concentration, neuronal swelling and ECS shrinkage, elevated extracellular glutamate duration). The experimental ranges are defined by the 25th percentile (Q1, splits off the lowest 25% of the data from the highest 75%) and the 75th percentile (Q3, splits off the highest 25% of the data from the lowest 75%) of the mean values reported in the experimental studies. The computational ranges are defined by the 25th and 75th percentiles of the values from model A, B, and C. To indicate variability outside the 25th and 75th percentiles, we define minimum and maximum values (the min/max-range) by the lowest and highest data points still within 1.5 IQR (where IQR = Q3 – Q1) of the lower and upper quartiles, respectively. Values outside the min/max-range are taken to be outliers. We qualitatively classify the match between computational and experimental results as follows: *in good agreement* if the computational and experimental intervals overlap, *overlap in range* if the min/max-ranges overlap but the intervals do not overlap, and *not in agreement* if the min/max-ranges do not overlap.

### 2.7 Code Accessibility

The code/software described in the paper is freely available online at: [https://bitbucket.org/adaje/supplementary-code-validating-a-computational-framework-for/src/master/]. The simulations were run on a Lenovo ThinkPad Carbon X1 11th Gen 2.80GHz × 8 Intel Core i7-1165G7 CPU with Ubuntu 20.04 using FEniCS 2019.1.0 without parallelization.

## 3 Results

### 3.1 Excitatory fluxes trigger wave of depolarization, ionic changes and swelling

In the wildtype computational model (A), the excitatory fluxes trigger a wave of neuronal and glial depolarization, changes in ionic concentrations and cellular swelling spreading through the tissue domain (Figure 2). We observe a depolarization of the neuronal and glial potentials from −68.5 mV to −15.5 mV and −82.0 mV to −38.9 mV, respectively, accompanied by a DC shift with an amplitude of 11.0 mV, with a duration of 38 s (Figure 2C, G). The neuronal depolarization wave is followed by an increase in the concentrations of extracellular K^+^ of 76.4 mM (Figure 2B, F), and glutamate of 1.38 mM (Figure 2A, E), and decreases in extracellular Na^+^ and Cl^−^ concentrations (Figure 2B, F). The increased levels of extracellular K^+^ and glutamate persist for 26 s and 20 s, respectively, whereas the drop in extracellular Cl^−^ lasts for 84 s. In response to the ionic shifts, both the neurons and the glial cells swell: we observe a neuronal swelling of 11.7% and a glial swelling of 7.13%; the extracellular space shrinks correspondingly. We observe altered volume fractions for 103–135 s (Table 2). Finally, the wave front has reached 5.25 mm after 60 seconds, and we observe a mean wave propagation speed of 5.84 mm/min. We note that the wave speeds *v*_*i*_ used to calculate the mean wave propagation speed, varies between 5.78 and 5.85 mm/min.

**Table 2.** Summary of computational quantities of interest for different models (A, B, C, D, E). Numerical errors are less than 1.5%.

| Quantity of interest | A | B | C | D | E |
| --- | --- | --- | --- | --- | --- |
| Mean wave propagation speed (mm/min) | 5.84 | 5.52 | 3.54 | 5.83 | 5.63 |
| DC shift (mV) | 11.02 | 3.80 | 9.37 | 11.02 | 8.98 |
| DC shift duration (s) | 32 | 50 | 69 | 32 | 42 |
| Neuronal swelling (%) | 11.69 | 11.67 | 6.90 | 14.93 | 12.16 |
| Glial swelling (%) | 7.13 | 7.05 | 4.05 | 0 | 4.37 |
| ECS shrinkage (%) | 39.79 | 39.56 | 9.37 | 37.33 | 36.81 |
| Neuronal swelling duration (s) | 109 | 111 | 178 | 115 | 103 |
| Glial swelling duration (s) | 103 | 101 | 177 | 0 | 143 |
| ECS shrinkage duration (s) | 135 | 135 | 179 | 136 | 131 |
| ECS K<sup>+</sup> elevation (mM) | 76.39 | 76.46 | 63.52 | 75.39 | 77.47 |
| ECS Glu elevation (mM) | 1.38 | 1.38 | 0.57 | 1.37 | 1.42 |
| ECS Cl<sup>-</sup> elevation (mM) | 21.12 | 20.88 | 9.72 | 23.12 | 18.63 |
| ECS K<sup>+</sup> elevation duration (s) | 26 | 26 | 49 | 26 | 27 |
| ECS Glu elevation duration (s) | 20 | 19 | 25 | 20 | 20 |
| ECS Cl<sup>-</sup> elevation duration (s) | 84 | 84 | 162 | 87 | 90 |
| Neuronal membrane potential (mV) | 63.36 | 63.38 | 57.38 | 63.32 | 63.58 |
| Glial membrane potential (mV) | 55.14 | 55.77 | 48.58 | 54.24 | 41.00 |
| Neuronal membrane potential duration (s) | 79 | 79 | 142 | 83 | 76 |
| Glial membrane potential duration (s) | 45 | 40 | 78 | 42 | 196 |

**Figure 2.**
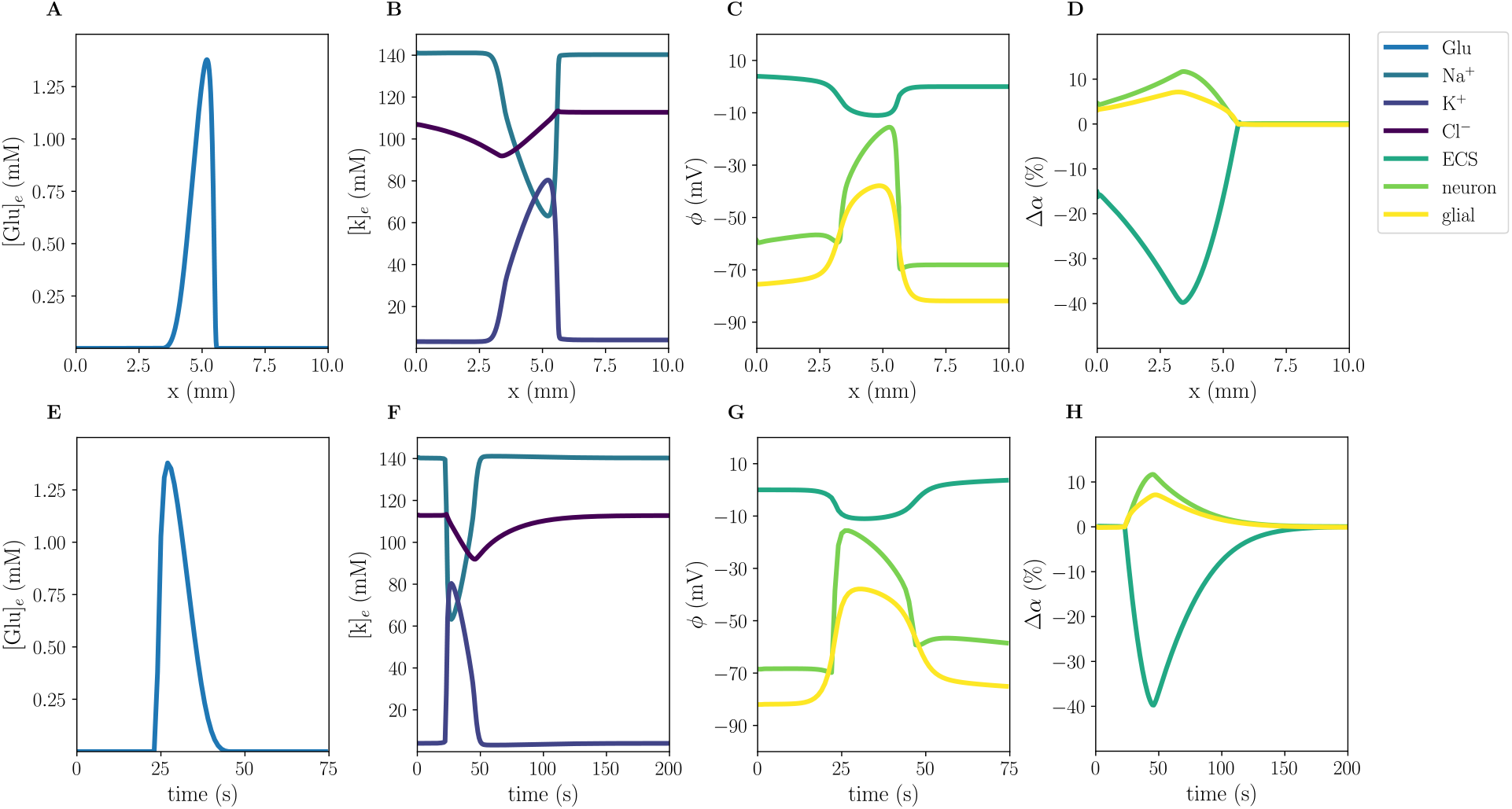
Simulated CSD wave in the wild type model (A) triggered by excitatory fluxes. The upper panels display a snapshot of ECS glutamate (**A**), ECS ion concentrations (**B**), potentials (**C**), and change in volume fractions (**D**) at 60 s. The lower panels display time evolution of ECS glutamate (**E**), ECS ion concentrations (**F**), potentials (**G**) and change in volume fractions (**H**) at *x* = 2.0mm.

### 3.2 Different computational CSD and SD triggers give comparable wave characteristics

CSD and SD can be triggered by different mechanisms experimentally, and we find that the same holds computationally. All three triggering mechanisms considered here – excitatory fluxes, topical application of K^+^ and disabled Na^+^/K^+^-ATPase (see Methods – induce a propagating SD wave with nearly identical wave characteristics including a mean wave propagation speed of 5.84 mm/min, 5.82 mm/min, and 5.83 mm/min, respectively.

### 3.3 Reduced glial intercellular diffusion reduces DC shift but maintains membrane potentials

The glial gap junction factor *χ*_*g*_ modulates the intercellular diffusion through the astrocytic networks. Reducing the effective intercellular diffusion (model B) does not lead to substantial changes in the CSD wave speed, ionic concentration changes, or cellular swelling (Table 2 model A vs B, Supplementary Figure 6C, G): model B gives a 5% reduction in the mean wave propagation speed (to 5.52 mm/min), and less than 2% in the other (ionic concentration or cellular swelling) quantities of interest. On the other hand, the DC shifts notably differ: the DC shift amplitude of model B (3.80 mV) is 66% smaller than in model A (11.02 mV). Yet, we observe that both the glial and neuronal potentials depolarize more in model B compared to model A, and thus the membrane potentials do not differ substantially between the two models.

### 3.4 Reduced membrane area-to-volume reduces CSD wave speed and amplitudes

Different values for the ratios of cell membrane area to unit tissue volume *γ* in brain tissue have been reported in the literature (Halnes et al. 2019, Kager et al. 2000). These parameters are thus uncertain and it is key to understand their effect on CSD wave characteristics. Reducing the membrane area-to-volume for the neurons *γ*_ne_ and glial cells *γ*_ge_ by respectively 71% and 68% (model C) substantially alters the CSD wave characteristics (Figure 3). In particular, the amplitudes of the ECS K^+^ and glutamate elevations are reduced by 17% and 59%, respectively. We observe that both neurons and glial cells swell less, and correspondingly the ECS shrinks less. Further, the amplitudes of the neuronal and the glial membrane potentials are respectively 11% and 12% smaller in the model C than in the model A (Table 2). Remarkably, the reduced membrane area-to-volume substantially slows down the CSD wave: the wave speed is 39% slower in WT-B (3.54 mm/min) compared to model A (5.84 mm/min) (Table 2).

**Figure 3.**
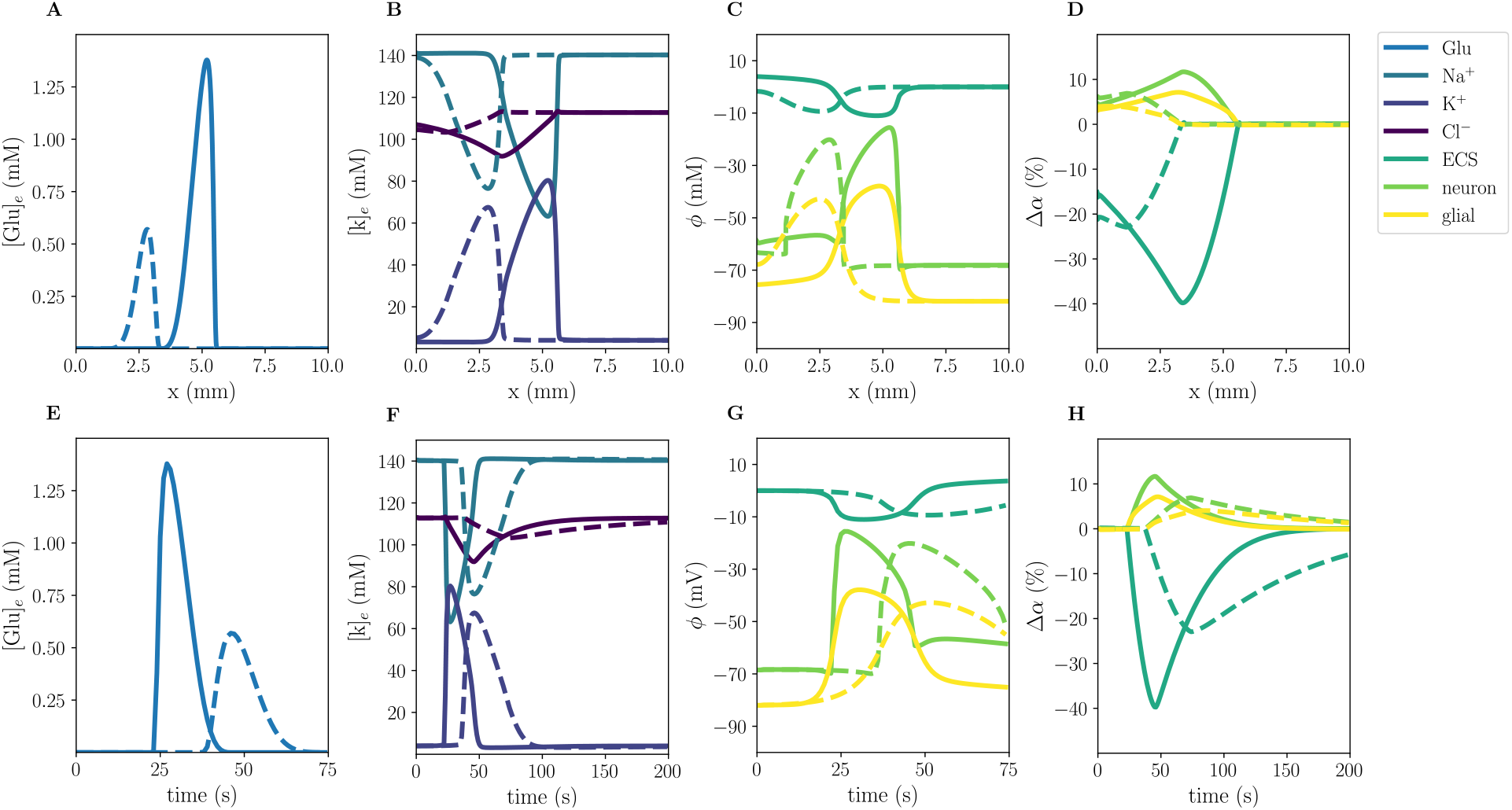
Comparison of model A (solid) and model C (stippled) CSD wave. The upper panels display snapshots of ECS glutamate (**A**), ECS ion concentrations (**B**), potentials (**C**), and change in volume fractions (**D**) at 60 s. The lower panels display time evolution of ECS glutamate (**E**), ECS ion concentrations (**F**), potentials (**G**) and change in volume fractions (**H**) evaluated at *x* = 2.0 mm.

### 3.5 Computational versus experimental quantities of interest in wild type mice

To evaluate the computational predictions, we compare our simulation results from the wild type models (A, B, C) with findings from a selection of experimental studies (summarized in Table 3). For the comparison, we define intervals of computational and experimental values for each of the relevant quantities of interest (Figure 4). The experimental ranges are defined by the 25th percentile and the 75th percentile of the mean values reported in the experimental studies. The computational ranges are defined by the 25th and 75th percentiles of the values from model A, B, and C. To indicate variability outside the 25th and 75th percentiles, we also define minimum and maximum values (the min/max-range, see Section 2.6 for details). Values outside the min/max-range are taken to be outliers.

**Table 3.**
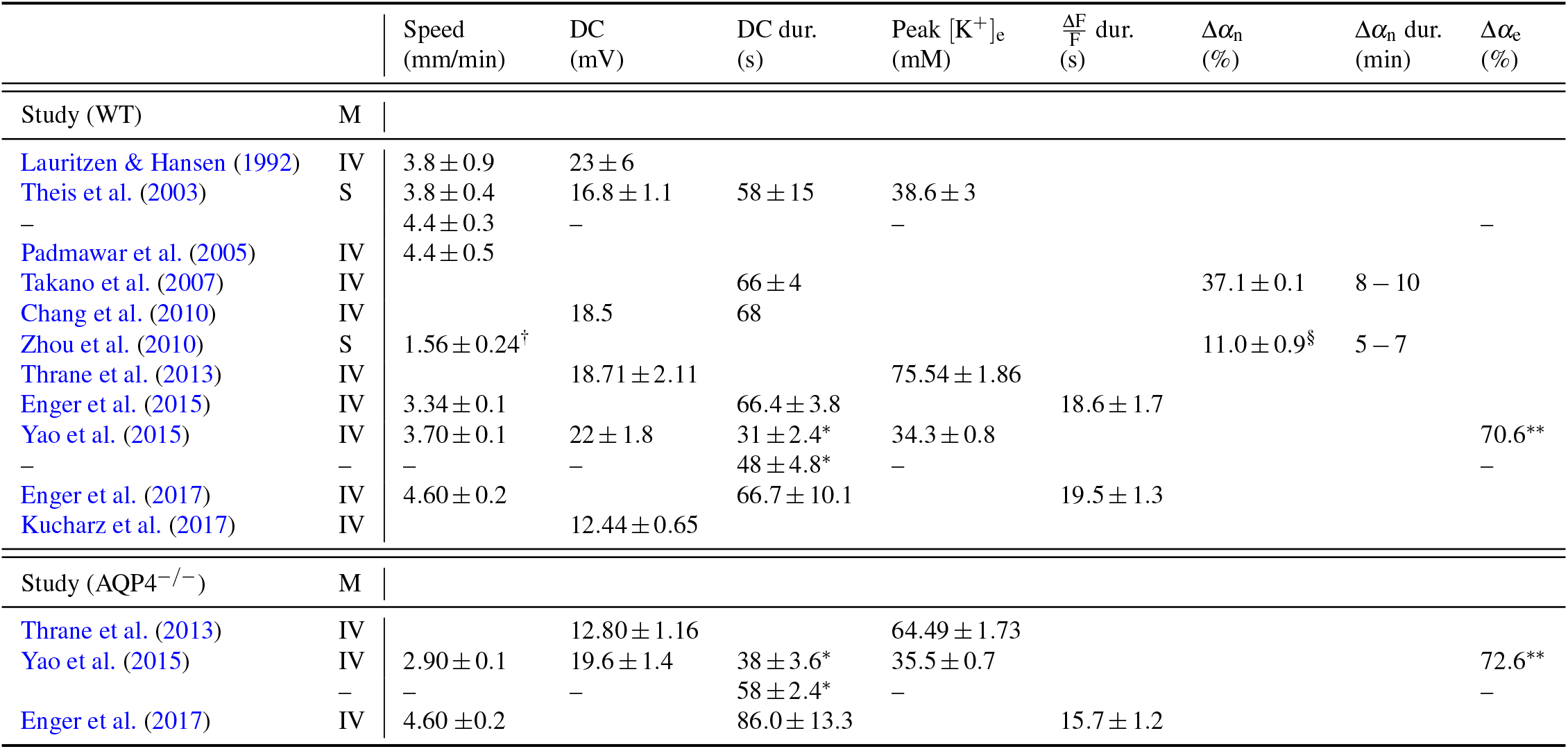
Summary and overview of experimentally reported propagation speeds, DC shift amplitudes (DC) and their duration (DC dur.), peak in extracellular K^+^ levels (Peak [K^+^]_e_), duration of increased relative changes in mean fluorescence 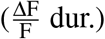, alteration in neuronal volume fractions (Δ*α*_n_) and their duration (Δ*α*_n_ dur.), and alteration in the extracellular space volume fractions (Δ*α*_e_) during CSD from a selection of studies in either wild-type mice (Study (WT)) or AQP4 knockout mice (Study (AQP4^−*/*−^)) measured (M) *in vivo* (IV) or in slices (S). ^*^ duration measured at half-maximum amplitude. ^†^ speed reported in experiments with TTX; indicated to be 50% of the CSD wave speed without TTX (Müller & Somjen 2000). ^§^ deviation from baseline of high [K^+^]_e_ perfusion. ^**^ baseline ECS volume fraction differs between WT mice (0.18) and AQP4^−*/*−^ mice (0.23).

| | | Speed<br>(mm/min) | DC<br>(mV) | DC dur.<br>(s) | Peak $[K^+]_e$<br>(mM) | $\frac{\Delta F}{F}$ dur.<br>(s) | $\Delta\alpha_n$<br>(%) | $\Delta\alpha_n$ dur.<br>(min) | $\Delta\alpha_e$<br>(%) |
| --- | --- | --- | --- | --- | --- | --- | --- | --- | --- |
| Study (WT) | M |  |  |  |  |  |  |  |  |
| Lauritzen & Hansen (1992) | IV | $3.8 \pm 0.9$ | $23 \pm 6$ | | | | | | |
| Theis et al. (2003) | S | $3.8 \pm 0.4$ | $16.8 \pm 1.1$ | $58 \pm 15$ | $38.6 \pm 3$ | | | | |
| – | | $4.4 \pm 0.3$ | – | | – | | | | – |
| Padmawar et al. (2005) | IV | $4.4 \pm 0.5$ | | | | | | | |
| Takano et al. (2007) | IV | | | $66 \pm 4$ | | | $37.1 \pm 0.1$ | 8 – 10 | |
| Chang et al. (2010) | IV |  | 18.5 | 68 |  |  |  |  |  |
| Zhou et al. (2010) | S | $1.56 \pm 0.24^\dagger$ | | | | | $11.0 \pm 0.9^\S$ | 5 – 7 | |
| Thrane et al. (2013) | IV | | $18.71 \pm 2.11$ | | $75.54 \pm 1.86$ | | | | |
| Enger et al. (2015) | IV | $3.34 \pm 0.1$ | | $66.4 \pm 3.8$ | | $18.6 \pm 1.7$ | | | |
| Yao et al. (2015) | IV | $3.70 \pm 0.1$ | $22 \pm 1.8$ | $31 \pm 2.4^*$ | $34.3 \pm 0.8$ | | | | $70.6^{**}$ |
| – | – | – | – | $48 \pm 4.8^*$ | – | | | | – |
| Enger et al. (2017) | IV | $4.60 \pm 0.2$ | | $66.7 \pm 10.1$ | | $19.5 \pm 1.3$ | | | |
| Kucharz et al. (2017) | IV | | $12.44 \pm 0.65$ | | | | | | |
| Study (AQP4<sup>-/-</sup>) | M |  |  |  |  |  |  |  |  |
| Thrane et al. (2013) | IV | | $12.80 \pm 1.16$ | | $64.49 \pm 1.73$ | | | | |
| Yao et al. (2015) | IV | $2.90 \pm 0.1$ | $19.6 \pm 1.4$ | $38 \pm 3.6^*$ | $35.5 \pm 0.7$ | | | | $72.6^{**}$ |
| – | – | – | – | $58 \pm 2.4^*$ | – | | | | – |
| Enger et al. (2017) | IV | $4.60 \pm 0.2$ | | $86.0 \pm 13.3$ | | $15.7 \pm 1.2$ | | | |

**Figure 4.**
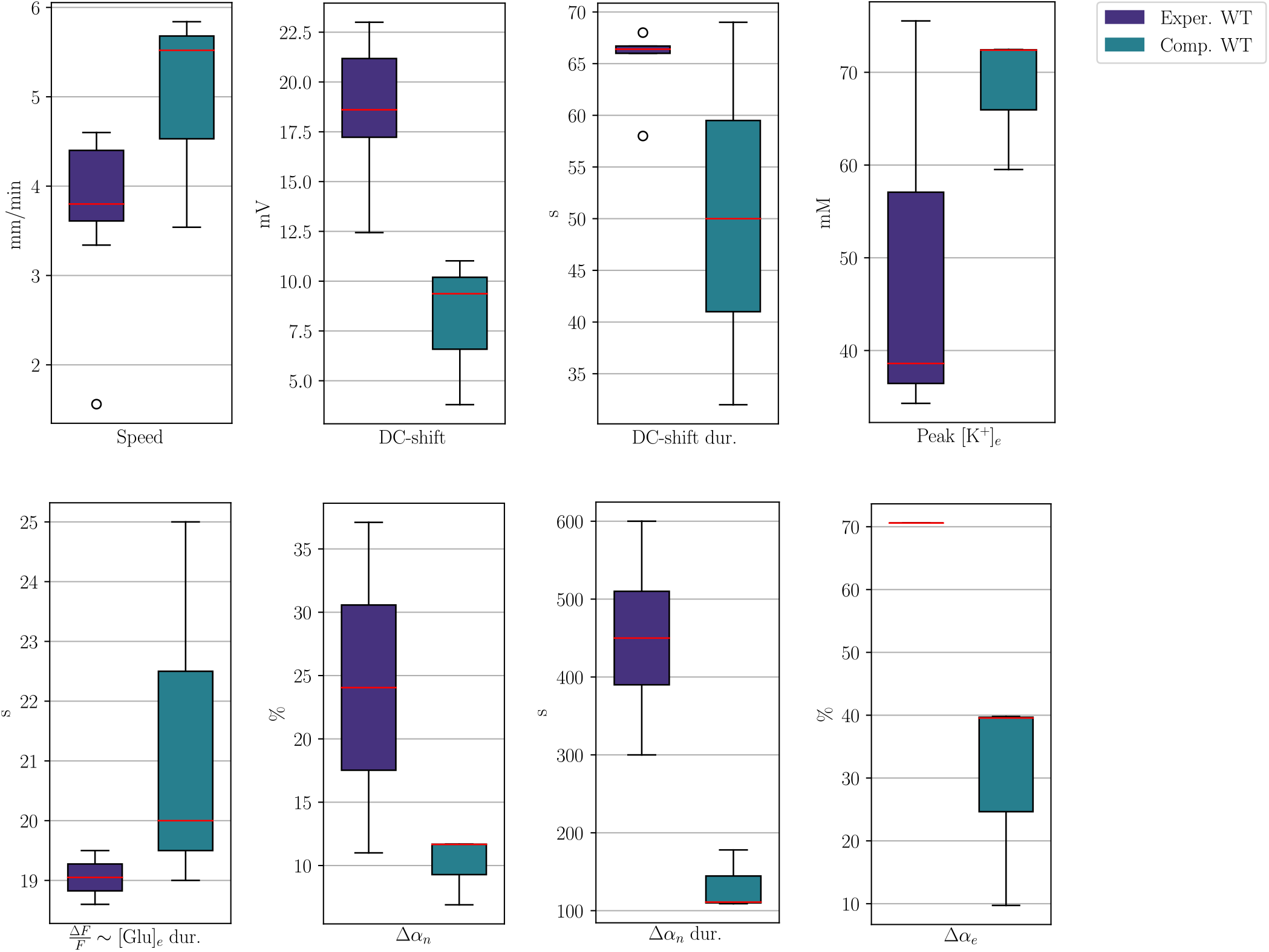
Comparison of intervals (boxes) and min/max-ranges (whiskers) for experimentally and computationally measured values for relevant quantities in wild type mice. Red lines and dots indicate median values and outliers, respectively.

CSD wave propagation speeds in wild type mice are reported in the min/max-range [3.34, 4.6] mm/min (Table 3), whereas the computational wild type models give max/min mean wave propagation speeds between 3.54 and 5.84 mm/min (Table 2). For the wave speed, there is thus overlap in the ranges of the computational and experimental results (Figure 4). As for the DC shift, the experimental values are in the min/max range [12.44, 23] mV, whereas we observe DC shifts within a min/max range of [6.59, 10.20] mV in the numerical simulations. We note that the lowest computational DC shift of 3.8 mV (model B) likely is an underestimate, as we expect some diffusion through the astrocytic networks. We thus find that the computational DC shifts are not in agreement with the experimental values. Regarding the duration of the DC shift, the experimental min/max-range ([66, 66.7] s, excluding measurements at half maximum amplitude) thus overlap with the computational min/max-range ([32, 69] s, Figure 4).

The computational values for the peak in extracellular K^+^ concentration overlap with the experimental reports (min/max-range of [59.52, 72.46] v.s. [38.6, 75.54] mM). Experimentally, elevated glutamate levels are typically indicated via the relative change in mean fluorescence (ΔF*/*F) over time (Enger et al. 2017, 2015). Comparing their duration, we find that elevation of extracellular glutamate in our computational models last for 19 to 25s, whereas the duration of the increase in ΔF*/*F has been reported experimentally in a min/max-range of [18.6, 19.5] s. The ranges of elevated extracellular glutamate duration thus overlap.

Neuronal swelling is reported in the min/max-range [11, 37.1]% in the collection of experimental studies, with a duration of minimum 300 seconds and maximum 600 seconds. We observe a neuronal swelling in the min/max-range [6.9, 11.69]%, lasting for 109 s to 178 s. The range of computational neuronal swelling amplitudes thus overlap with the experimental range, whereas the computational durations do not agree with experimental reports. We also note that neither Zhou et al. (2010) nor Takano et al. (2007) find that astrocytes swell significantly during CSD. In contrast, our numerical findings show glial (astrocytic) swelling within a range of [4.05, 7.13]%. Regarding extracellular shrinkage, Yao et al. (2015) report a 70.6% reduction of the ECS volume. The computational values are not in agreement: predicting a reduction in the extracellular volume fraction in a min/max-range of [9.37, 39.79]%.

### 3.6 Reducing the glial water permeability affects cellular swelling

When setting the glial water permeability to zero, we naturally observe no glial swelling, while the neuronal swelling increases by 27.7% compared to the baseline and the resulting shrinkage of the extracellular space is 6.2% smaller (model D, Table 2 and Supplementary Figure 7). The remaining CSD wave characteristics, including the CSD wave propagation speed, do not change notably.

Experimentally, Thrane et al. (2013), Yao et al. (2015), and Enger et al. (2017) have studied CSD in AQP4 knockout mice. Enger et al. (2017) observe no difference in the CSD wave propagation speeds between the WT and AQP4 knockouts (4.6± 0.2 mm/min). This is in contrast to the findings of Yao et al. (2015), who report that the CSD wave propagation speed is reduced by 22% in AQP4 knockouts. Yao et al. (2015) and Thrane et al. (2013) also report a small, but significant, reduction in the DC shift amplitude in AQP4 knockouts compared to wild type mice, whereas Enger et al. (2017) observe no significant difference in either the duration or the amplitude of the DC shift between AQP4^−*/*−^ and WT. Enger et al. (2017) additionally study extracellular glutamate elevations during CSD. They report a 20% reduction in the duration of glutamate elevation for AQP4 knockouts (18.6± 1.7 s in WT v.s. 15.7 ±1.2 s in AQP4 knockouts). Moreover, Thrane et al. (2013) report that the amplitude of elevated levels of extracellular K^+^ is significantly lower in AQP4^−*/*−^ than in WT mice. Our computational findings (modulo the duration of glutamate elevation) are thus in agreement with the experimental results of Enger et al. (2017).

### 3.7 Reduced K_ir_4.1 expression changes glial and DC shift dynamics

Reducing the K_ir_4.1 channel conductivity (model E) alters the CSD wave characteristics, inducing changes in the glial potential, glial swelling and DC shift (Table 2, Figure 5). In particular, we observe that the glial membrane potential amplitude drops from 55.14 to 41.00 when the K_ir_4.1 expression is reduced. Moreover, we observe a 39% reduction in glial swelling amplitude, a 4% increase in neuronal swelling amplitude and a corresponding decrease in the ECS shrinkage amplitude. Reducing K_ir_4.1 also results in a slightly higher ECS K^+^ amplitude (1.4%), a lower ECS glutamate amplitude (11.8%), and a slightly reduced wave speed (3.6%).

**Figure 5.**
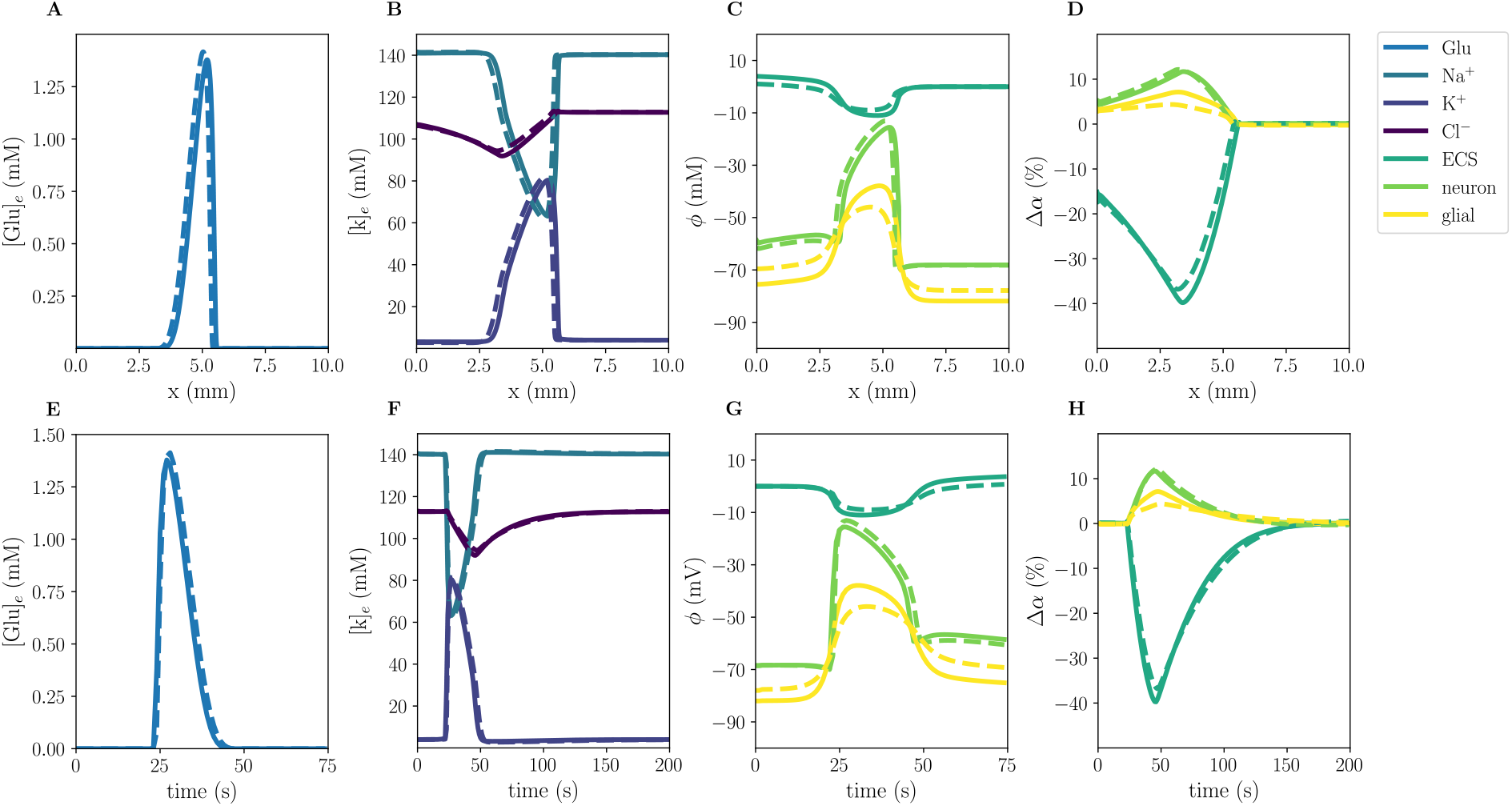
Comparison of model A (solid) and model E (stippled) CSD wave. The upper panels display snapshots of ECS glutamate (**A**), ECS ion concentrations (**B**), potentials (**C**), and change in volume fractions (**D**) at 60 s. The lower panels display time evolution of ECS glutamate (**E**), ECS ion concentrations (**F**), potentials (**G**) and change in volume fractions (**H**) evaluated at *x* = 2.0 mm.

## 4 Discussion

In this study, we have simulated CSD in computational models including multiple triggering mechanisms under variations in morphological properties, intercellular diffusivity, membrane water permeabilities, and channel conductance, and compared the computational findings with experimental reports. The range of wave speeds, DC shifts duration, peak extracellular K^+^ concentration, neuronal changes in volume fraction, and extracellular shrinkage obtained computationally all overlap with the experimentally reported ranges. Our findings show that intercellular glial diffusion strongly affects the DC shift, and that the ratio of cellular membrane area to tissue volume strongly affects the CSD wave speed. The computational model predicts that a reduced K_ir_4.1 expression will reduce the glial swelling and the depolarization of the glial membrane during CSD.

While several papers consider mathematical and computational modelling of CSD (Almeida et al. 2004, Florence et al. 2009, Kager et al. 2000, 2002, Mori 2015, O’Connell & Mori 2016, Shapiro 2001, Tuttle et al. 2019, Wei et al. 2014), few have performed an systematic comparison with experimental literature. The effects of varying glial gap junction strength and the K_ir_4.1 conductance on the DC shift and CSD wave speed has previously been computationally assessed by O’Connell & Mori (2016). They find that the glial gap junction coupling impacts the DC-shift, which is in agreement with our computational results. Further, they find that the glial cell either buffer or broadcast K^+^, depending on the values of the gap junction and K_ir_4.1 conductance. To our knowledge, there are no computational studies examining the effects of reduced glial water permeability or exploring the effect of membrane area-to-volume ratios during CSD.

There are several experimental studies on the role of AQP4 in CSD (Enger et al. 2017, Rosic et al. 2019, Thrane et al. 2013, Yao et al. 2015). Astroglial transmembrane water transport may also occur via other membrane mechanisms (e.g. EAAT, NKCC, VRAC) (MacAulay et al. 2004, Pedersen et al. 2016), and the individual contributions to the total transmembrane water permeability of these different mechanisms are debated. Instead of targeting specific water mechanisms, we have here studied the effect of reducing the total glial transmembrane water permeability. When eliminating the glial membrane water transport entirely (model D), we find, as expected, that the glial cells do not swell at all, whereas the neuronal swelling increases in comparison with the baseline model (model A). We do not find any difference in CSD wave speed nor in any other computational quantity of interest between these two models. We note that the experimental literature is inconclusive with regard to this point. Our computational findings are largely in agreement with the experimental findings of Enger et al. (Enger et al. 2017) – knocking out AQP4 does not substantially alter the CSD wave characteristics. In contrast Yao et al. (2015) and Thrane et al. (2013) report significant differences in their AQP4^−*/*−^ mice. It is however clear that AQP4 knockout mice may have additional morphological differences, e.g. related to the extracellular space volume: Yao et al. (2008, 2015) report a ∼30% larger ECS baseline volume in AQP4^−*/*−^ than in wildtype mice. Our findings suggest that if a reduction in CSD wave speed in AQP4^−*/*−^ is is present, it originates from other factors than the water permeability of the glial membrane. In particular, the ratio between glial and neuronal membrane area and tissue volume was found to be an important factor for the CSD wave speed.

Diffusion of extracellular K^+^ has been hypothesized to be an underlying mechanisms in CSD wave propagation (Grafstein 1956, Obrenovitch & Zilkha 1995, Pietrobon & Moskowitz 2014). Local elevations in [K^+^]_e_ will, via diffusion, increase the levels of [K^+^]_e_ in neighboring regions, activating voltage-gated and/or [K^+^]_e_-dependent channels which leads to a further depolarization of the membrane and thus further release of K^+^ into the extracellular space. Our models predict extracellular K^+^ amplitudes and wave speeds that are in line with studies performed without the dampening effects of anesthesia; thus we conjecture that these observations in our computational results may be connected. Indeed, in the model with reduced membrane area-to-volume, we observe lower extracellular K^+^ amplitudes and lower wave speeds.

Spatial K^+^ buffering by astrocytes is hypothesized to be a key mechanism in controlling extracellular K^+^ levels. Astrocytes buffer and redistribute extracellular K^+^ via the astrocytic networks by transferring K^+^ ions from regions with high K^+^ concentration to regions with lower K^+^ levels (Kofuji & Newman 2004, Orkand et al. 1966). There are open questions related to the role of spatial K^+^ buffering in CSD. Spatial buffering may prevent buildup of extracellular K^+^, or it could potentially increase the CSD wave speed through K^+^ distribution in the direction of the wave propagation. The influx of K^+^ during spatial buffering is mainly mediated by the inward rectifying K^+^ channel K_ir_4.1 (Ohno 2018, Rasmussen et al. 2020). In general, studying the role of K_ir_4.1 channels *in vivo* is challenging: the lack of K_ir_4.1 has been reported to cause hyper-excitable neurons, epileptic seizures, and premature death (within 3-4 weeks) in mice (Djukic et al. 2007). Modelling K_ir_4.1 knockouts computationally could potentially aid in investigating the role of K_ir_4.1. Our findings show that a reduced K_ir_4.1 expression will reduce glial swelling and the depolarization of the glial membrane during CSD. We also observe a slightly reduced CSD wave propagation speed. Further, glial buffering currents have been suggested as an important mechanism behind DC potentials in brain tissue (Herreras & Makarova 2020, Sætra et al. 2021). This is in line with our observation that a reduction in glial intercellular diffusion reduces the DC shift: reducing glial intercellular diffusion will effectively remove the glial buffering currents.

In terms of limitations, the model framework is founded on an homogenized representation of the tissue: the extracellular space, the cell membrane and the intracellular space are assumed to exist everywhere in the computational domain. For the spatial and temporal scales involved in the propagation of CSD, such a representation seems appropriate. Next, our simulations are based on a one-dimensional computational domain. As CSD waves spread through the whole depth of the cortical tissue, waves in two or three dimensions would be a more accurate representation. We would expect the computational CSD wave speeds to be reduced in centre-initiated two-dimensional simulations. We also note that experimental studies often measure relative fluorescence intensity instead of glutamate concentrations directly, while substantial uncertainty is associated with those that do, e.g. Fabricius et al. (1993), Scheller et al. (2000). We have therefore not compared this model quantity with experimental data.

The astroglial membrane expresses a variety of voltage-dependent and leak K^+^ channels (Olsen 2012), while our mathematical model only accounts for K_ir_4.1. The lack of other K^+^ model mechanisms limits the computational range of K_ir_4.1 permeabilities: reductions further than 70% leads to numerical instabilities. We hypothesize that a further reduction of the K_ir_4.1 expression could be obtained by including other glial K^+^ membrane channels (e.g. K^+^ leak channels). For further computational studies of K_ir_4.1 dynamics, it would indeed be advantageous to reduce the K_ir_4.1 expression to the point where the resting glial membrane potential is around −58 mV, as observed *in vivo* (Chever et al. 2010). Differences in CSD DC shift duration between different sites have been noted in experiments (Theis et al. 2003, Yao et al. 2015). In our computational model, due to the homogeneity of the material parameters, we expect (and observe) little spatial variation in the computed quantities of interest. Including heterogeneity and/or anisotropy in the model parameters would allow for studying how local tissue properties effect CSD dynamics.

Validating complex computational models with numerous state variables and model parameters against experimental findings with considerable variability is challenging. Although the methodology suggested here captures variability between the experimental studies, it fails in capturing the uncertainty (i.e. the standard deviation) within each individual study. Conducting rigorous meta-analysis by combining data from the individual experiments, weighted by e.g. the studies quality, size and/or other factors, to a pooled estimate for the experimental quantities of interest would be advantageous. However, the methodological differences (e.g. recording techniques, recording sites, *in vivo*, awake or anesthetized, v.s. *in vitro*) between the experimental studies makes meta-analysis difficult (Greco et al. 2013). The computational model considered within this work is complex with numerous model parameters, many of which are difficult to measure experimentally, giving rise to considerable uncertainty. A thorough uncertainty and sensitivity analysis of the computational CSD model would be interesting for future work. Importantly, the model constitutes a general yet detailed framework for capturing fundamental mechanisms of the interplay between ionic movement and volume control of the extracellular space. As such, applications of the framework are not limited to CSD.

We conclude that the Mori framework is a promising tool for predicting complex phenomena of ionic electrodiffusion and osmotic dynamics in brain tissue. Our findings indicate that the computational model predictions qualitatively matched well with experiments when applied to CSD. As a general framework for ionic dynamics and osmosis, applications are not limited to CSD. Mathematical modelling can be useful for isolating effects and e.g. pointing at potential confounding effects in AQP4 knockout mice and how they differ from wild type in various aspects.

## A Supplementary Methods

The mathematical model for CSD through cerebral tissue outlined below is introduced by Mori and coauthors (Mori 2015, O’Connell & Mori 2016, Tuttle et al. 2019). The tissue of interest is represented as a domain Ω∈ℝ^*d*^, with *d* = 1, 2, 3. Moreover, the tissue is composed of *N* interpenetrating compartments. We assume that *r* = *N* always denotes the ECS, and that other compartments communicates directly with the ECS only. Throughout this note, we will consider three compartments (*N* = 3), namely a neuron compartment (r=1, denoted by n), a glial compartment (r=2, denoted by g) and an extracellular compartment (r=3, denoted by e). Consider time *t* ∈ (0, *T*].

### A.1 Governing equations

We will consider the following system of coupled, time-dependent, nonlinear PDEs. Find the *volume fraction α*_r_ : Ω ×(0, *T*] → ℝ such that for each *t* ∈ (0, *T*]:

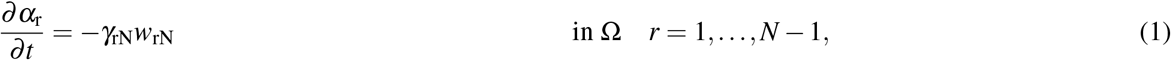

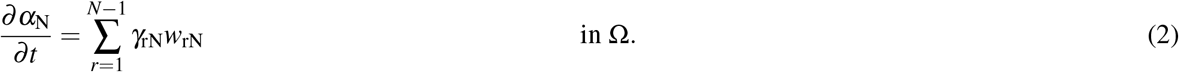

The *transmembrane water flux w*_rN_ between compartment *r* and the ECS is driven by osmotic and oncotic pressure, and will be discussed further in Section A.2. The coefficient *γ*_rN_ (m^−1^) represents the area of cell membrane between compartment *r* and the ECS per unit volume of tissue. By definition we have that

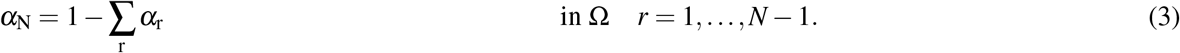

For each ion species *k* ∈ *K*, find the *ion concentrations* [*k*]_r_ : Ω ×(0, *T*] → ℕ (mol/m^3^) and the *electrical potentials ϕ*_r_ : Ω ×(0, *T*] → ℝ (V) such that for each *t* ∈ (0, *T*]:

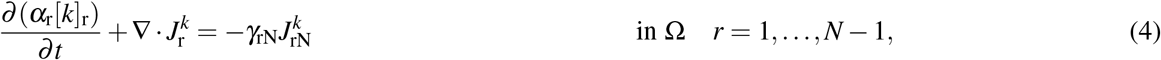

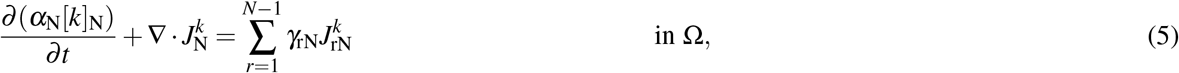

where 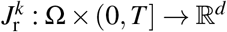 is the *ion flux density* (mol/(m^2^s)) for each ion species *k*. The *transmembrane ion flux density* (mol/(m^2^s)) 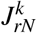 for each ion species *k* is subject to modelling, and will be discussed further in Section A.2. Note that (4)–(5) follow from first principles and express conservation of ion concentrations for the bulk of each region. Moreover, we assume that the ion flux densities (and *a–fortiori* the ion concentrations and electrical potentials) satisfy:

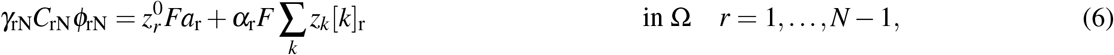

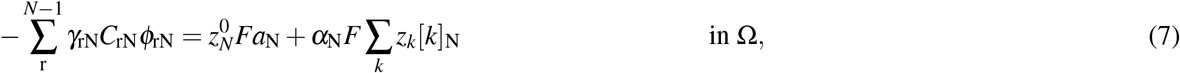

where *z*^*k*^ is the *valence* of ion species *k*, and *C*_rN_ is the *membrane capacitance* (F/m^2^) of the membrane between compartment *r* and the ECS. In this paper, we assume that the ion flux densities can be expressed in terms of the ion concentrations, the electrical potentials, and the volume fractions as:

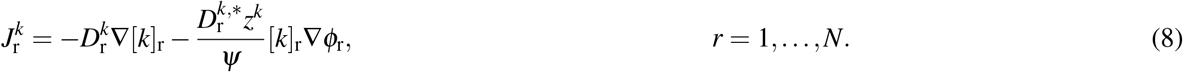

Here, the constant *ψ* = *RTF*^−1^ combines *Faraday’s constant F*, the *absolute temperature T*, and the *gas constant R*. Furthermore,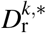 denotes the *effective diffusion coefficient* of ion species *k* in the region *r* and may be a given constant or e.g. a spatially varying scalar field dependent of *α*_r_. In particular, we let the effective diffusive coefficients 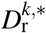 be:

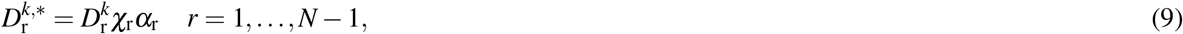

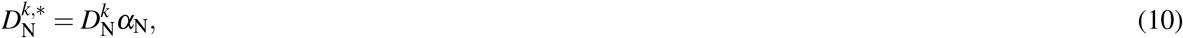

where 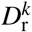 is the diffusion coefficient of ion species *k* in region *r* and *χ*_r_ represents the cellular gap junctions in region *r*. The ionic flux density, i.e. the flow rate of ions per unit area, is thus modelled as the sum of three terms: (i) the diffusive movement of ions due to ionic gradients 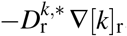, (ii) the ion concentrations that are transported via electrical potential gradients, i.e. the ion migration 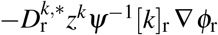 where 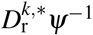 is the *electrochemical mobility* and (iii) the convective movement *α*_r_*u*_r_[*k*]_r_ of ions. Equations (1)–(7) with insertion of (8), thus constitutes a system of |*N*| |*K*| + 2 |*N*| + 1 equations for the |*N*|| *K*| + 2| *N* |+ 1 unknown scalar fields. Appropriate initial conditions, boundary conditions, and importantly membrane mechanisms must be set in order to close the system.

### A.2 Membrane mechanisms

The transmembrane water flux *w*_rN_ is driven by a combination of hydrostatic and osmosis pressure, and can thus be expressed as:

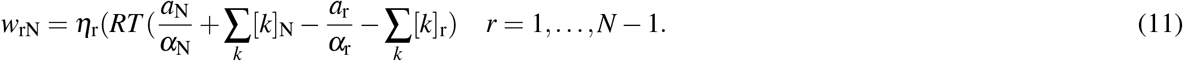

The intracellular and extracellular potentials across the membrane is coupled by letting *ϕ*_rN_ be the jump in the electrical potential across the membrane between compartment *r* and the ECS. Thus, the membrane potential can be expressed as the difference between the intracellular and extracellular potentials:

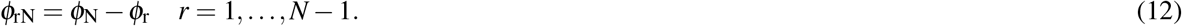

The leak currents (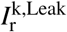,A/m^2^) of ion species *k* over the membrane between the intracellular and extracellular compartments are modeled as:

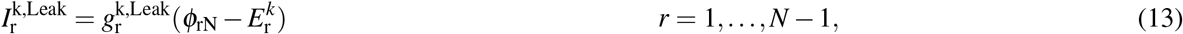

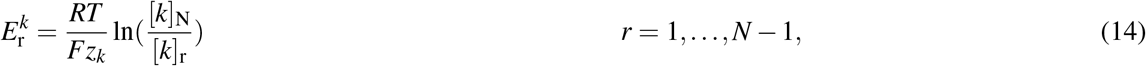

where 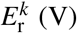 denotes the Nernst potential, whereas the leak conductances (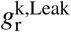,S/m^2^) are listed in Table 6. The glial cells are more inclined to absorb than to release K^+^. Therefore, the conductance for the K^+^ inward rectifier current (K_ir_4.1, A/m^2^) is modelled as an expression depending on the extracellular and intracellular ion concentrations for the glial cell. Following Steinberg et al. (2005), the K_ir_4.1 conductance (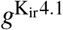,S/m^2^) is modelled as:

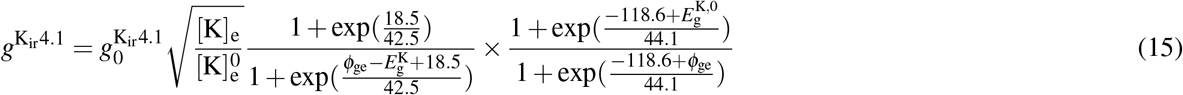

where 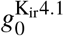 (S/m^2^) is the resting membrane conductance, and corresponds to the conductance when 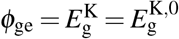 and 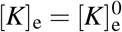. The value of 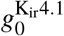 can be found in Table 6.

The voltage-dependent Na^+^ and K^+^ conductances are simulated using the classical Hodgkin-Huxley kinetic description. The current-voltage relation for the voltage-gated current (*I*^*k*,GHK^, A/m^2^) of ion species *k* over the neuron membrane is described by the Goldman-Hodgkin-Katz (GHK) current equation:

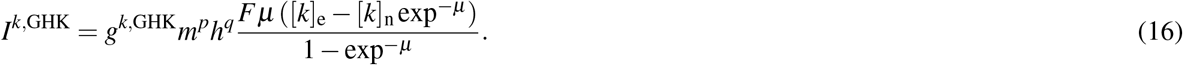

Here, 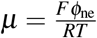 is a dimensionless value, while *g*^*k*,GHK^ (m/s) denotes the product of membrane permeability and conductance. The gating variables *m* and *h* describe the proportion of open voltage–gated ion channels, and are governed by the following ODEs:

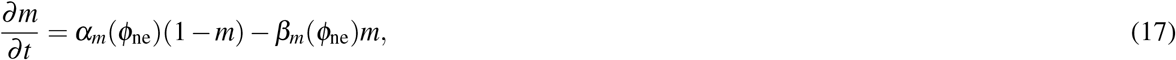

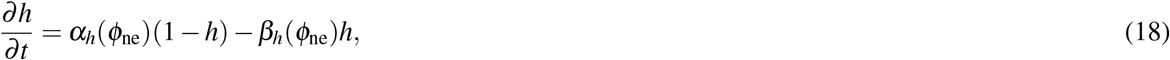

where the activation rate functions *α*_*m*_ : ℝ →ℝ and *β*_*m*_ : ℝ→ ℝ, and the inactivation rate functions *α*_*h*_ : ℝ→ ℝ and *β*_*h*_ : ℝ →ℝ, are specified for each current in Table 4. The gating variable for the NMDA receptor is determined by *y*, which is governed by the following ODEs:

**Table 4.**
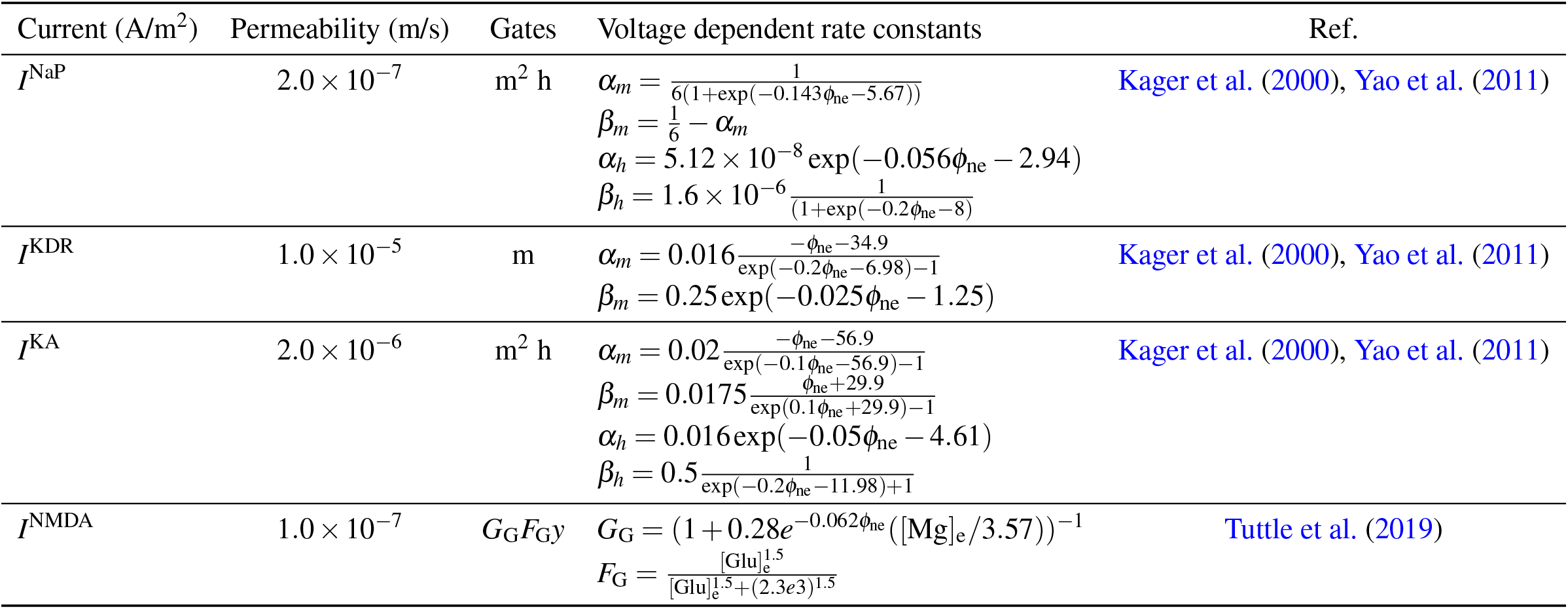
Permeability, gates and voltage dependent expressions for the activation rates (*α*) and the inactivation rates (*β*) for the persistent Na^+^ current (NaP), the K^+^ delayed rectifier current (KDR) and the transient K^+^ current (KA) in the neuron membrane and model for the NMDA receptor current.

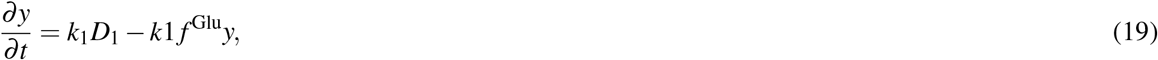

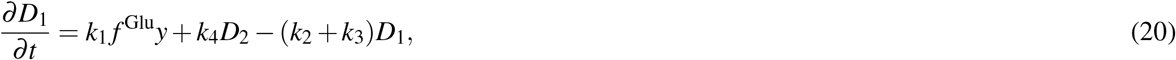

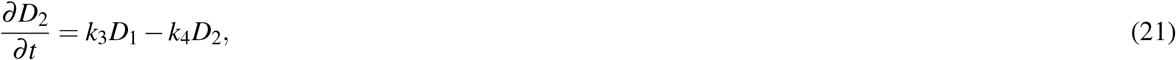

where *k*_1_, *k*_2_, *k*_3_ and *k*_4_ are transition rates between the states. We have included four voltage gated currents in the neuron membrane, namely *persistent Na*^+^ *current* (NaP), *K*^+^ *delayed rectifier current* (KDR), *the transient K*^+^ *current* (KA), and *N-methyl-D-aspartate* (NMDA). The Na^+^/K^+^-ATPase (ATP) occur in both the neuron and glial membrane. The pump exchanges 2 K^+^ ions for 3 Na^+^ ions, and the pump currents (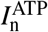 and 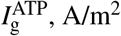 over the neuronal and glial membranes, respectively, are modelled as:

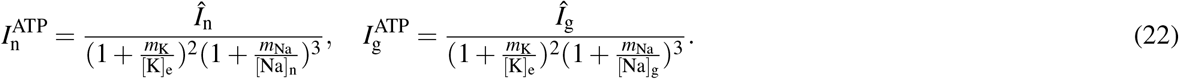

The Na^+^, K^+^, Cl^−^ co-transporters (NaKCl) occur in the glial membrane only, and the co-transporter current *I*^NaKCl^ (A/m^2^) over the glial membrane is modelled as:

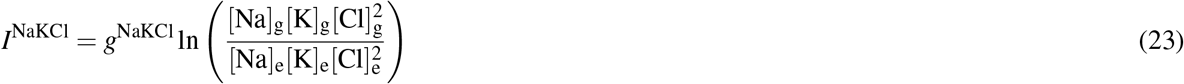

Glutamate exchange between the neurons, the glial cells and the extracellular space is modelled through the following expressions:

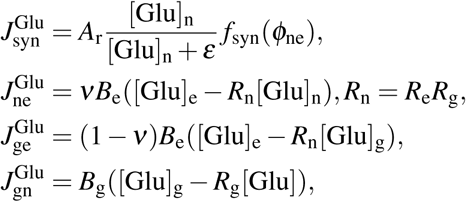

where *A*_r_ denotes the release rate of glutamate, *ε* denotes the saturation constant, *ν* denotes the reabsorption rate percentage, and *B*_e_ and *B*_g_ denote the decay and cycle rate, respectively. Moreover, *R*_e_ and *R*_g_ denote the ECS and glial fractions, respectively. Finally, we have that:

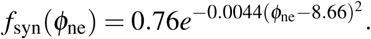

In summary, the total currents over the neuron and glial membrane is modelled by (24). In order to convert the currents (A/m^2^) to ion fluxes (mol/(m^2^s)), we multiply the currents by the valence *z*_*k*_ and divide by Faraday’s constant *F*:

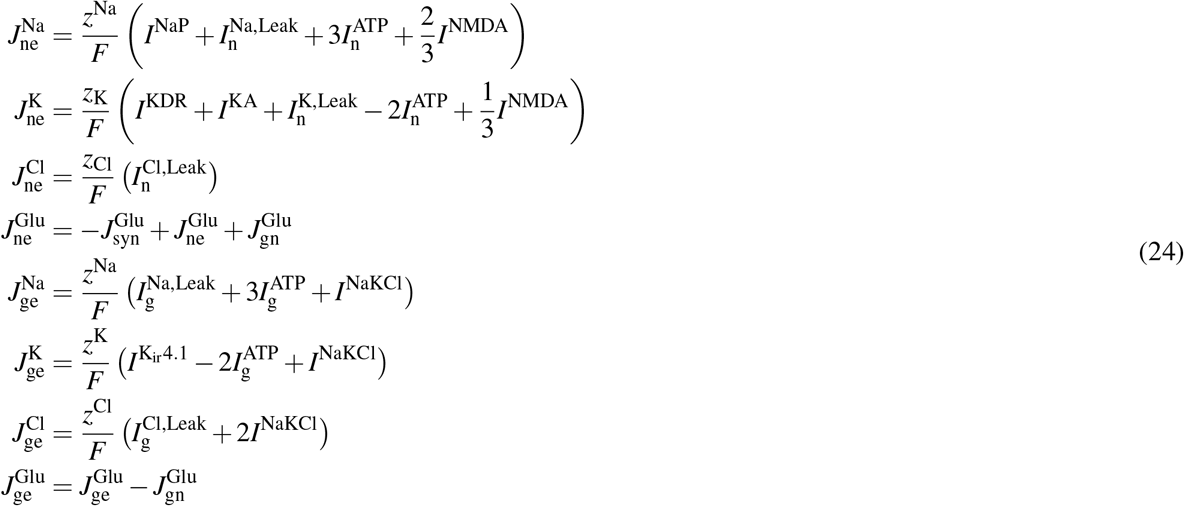

### A.3 Parameter values

The parameter values are as given in Table 5 (tissue parameters) and in Table 6 (membrane model parameters) unless otherwise specified.

**Table 5.**
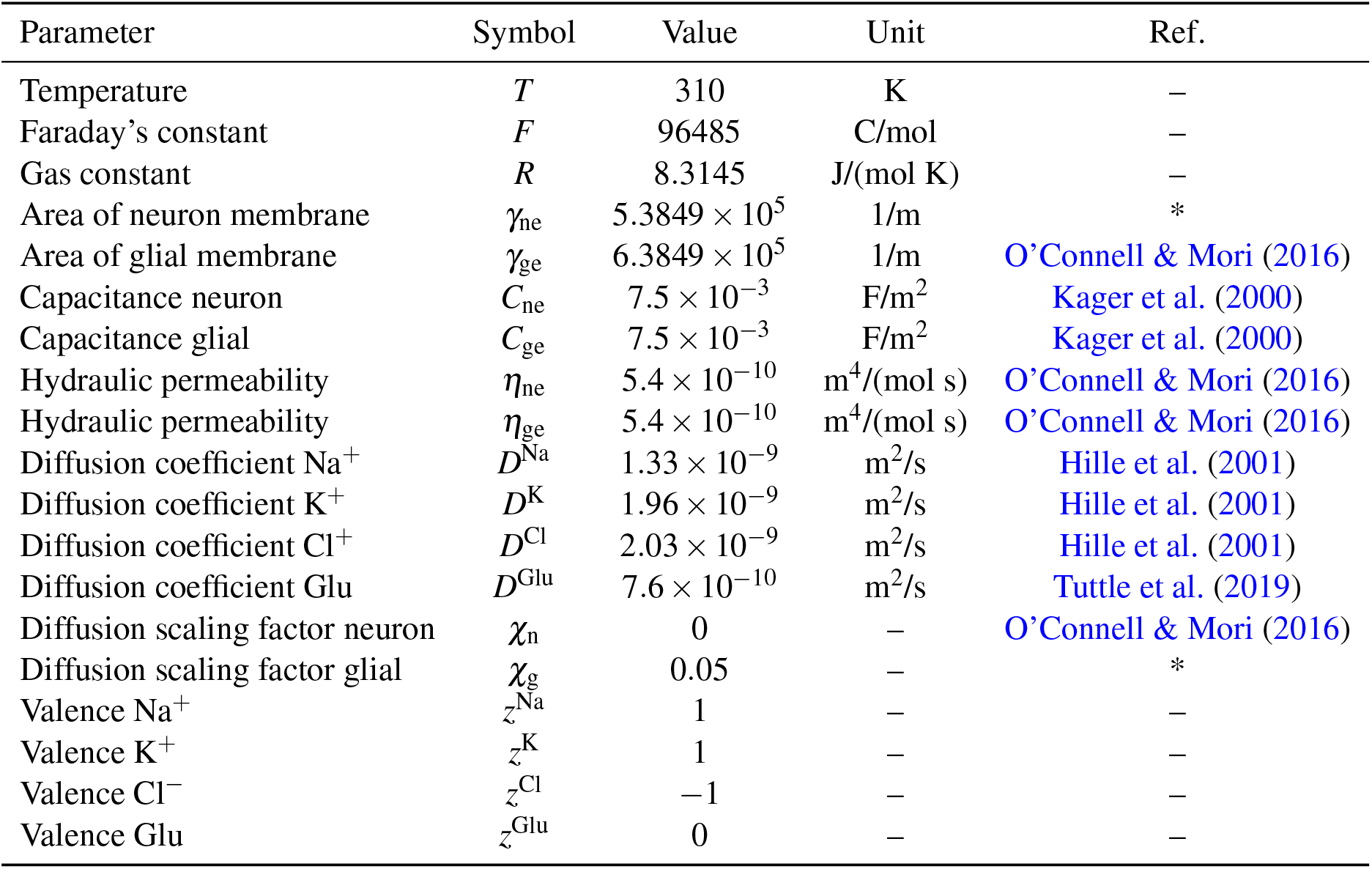
Tissue model parameters (default). We use SI base units, that is, Kelvin (K), Coulomb (C), mole (mol), meter (m), second (s), Joule (J) and Pascal (Pa). The hyphen (–) indicates that the parameter is dimensionless (Unit column), or that a standard value is used (Ref.column).

**Table 6.**
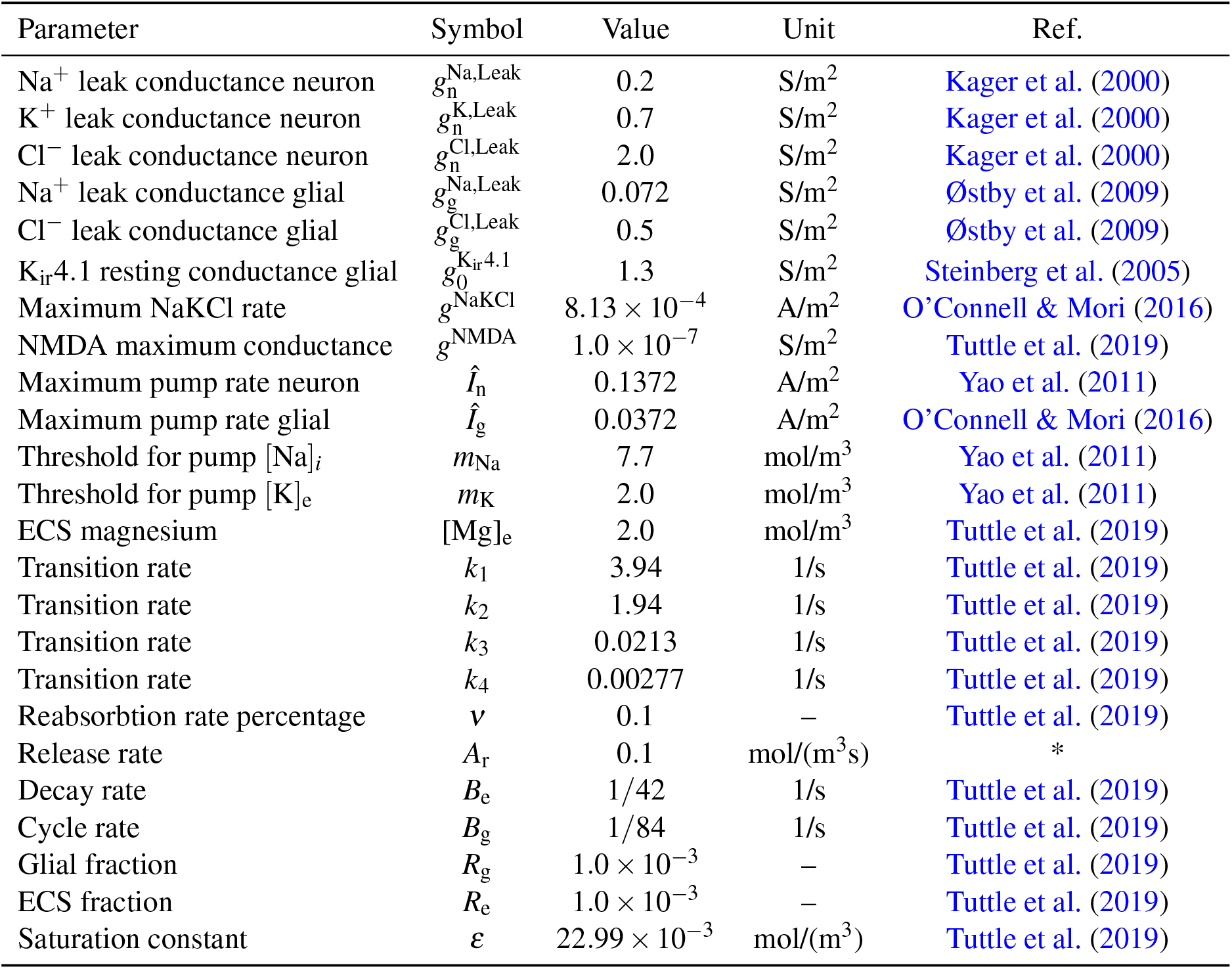
Membrane model parameters (default). We use SI base units; that is, meter (m), second (s), mole (mol), Siemens (S) and ampere (A).

### A.4 Boundary conditions

The boundary conditions will strongly depend on the problem of interest. If not otherwise stated, we assume that no ion flux leave at exterior boundary *∂* Ω, that is:

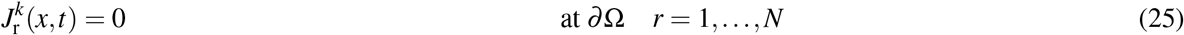

### A.5 Initial conditions

The initial conditions for the PDE system 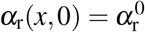, and 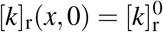 are set such that the system is is steady state without stimuli (Table 7). The initial conditions for the ODE system are set in the following manner:

**Table 7.** Initial values for state variables. We use SI base units; that is, meter (m), and mole (mol). The hyphen (–) indicates that the parameter is dimensionless (Unit column), or that a standard value is used (Ref. column), whereas * indicate that the value was calculated to obtain steady state.

| Parameter | Symbol | Value | Unit | Ref. |
| --- | --- | --- | --- | --- |
| neuron volume fraction | $\alpha_n^0$ | 0.5 | – | O’Connell & Mori (2016) |
| glial volume fraction | $\alpha_g^0$ | 0.3 | – | O’Connell & Mori (2016) |
| Na<sup>+</sup> concentration neuron | $[\text{Na}]_n^0$ | 9.3 | mol/m<sup>3</sup> | * |
| K<sup>+</sup> concentration neuron | $[\text{K}]_n^0$ | 130 | mol/m<sup>3</sup> | O’Connell & Mori (2016) |
| Cl<sup>–</sup> concentration neuron | $[\text{Cl}]_n^0$ | 8.7 | mol/m<sup>3</sup> | * |
| Glu concentration neuron | $[\text{Glu}]_n^0$ | 10 | mol/m<sup>3</sup> | Tuttle et al. (2019) |
| Na<sup>+</sup> concentration glial | $[\text{Na}]_g^0$ | 14 | mol/m<sup>3</sup> | * |
| K<sup>+</sup> concentration glial | $[\text{K}]_g^0$ | 130 | mol/m<sup>3</sup> | O’Connell & Mori (2016) |
| Cl<sup>–</sup> concentration glial | $[\text{Cl}]_g^0$ | 8.5 | mol/m<sup>3</sup> | * |
| Glu concentration glial | $[\text{Glu}]_g^0$ | $1.0 \times 10^{-3}$ | mol/m<sup>3</sup> | Tuttle et al. (2019) |
| Na<sup>+</sup> concentration ECS | $[\text{Na}]_e^0$ | 140 | mol/m<sup>3</sup> | O’Connell & Mori (2016) |
| K<sup>+</sup> concentration ECS | $[\text{K}]_e^0$ | 4 | mol/m<sup>3</sup> | O’Connell & Mori (2016) |
| Cl<sup>–</sup> concentration ECS | $[\text{Cl}]_e^0$ | 113 | mol/m<sup>3</sup> | O’Connell & Mori (2016) |
| Glu concentration ECS | $[\text{Glu}]_e^0$ | $1.0 \times 10^{-5}$ | mol/m<sup>3</sup> | Tuttle et al. (2019) |
| potential neuron | $\phi_n^0$ | –0.069 | V | * |
| potential glial | $\phi_g^0$ | –0.082 | V | * |
| potential ECS | $\phi_e^0$ | 0.0 | V | – |

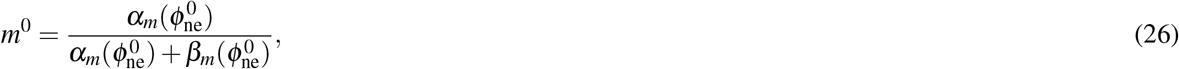

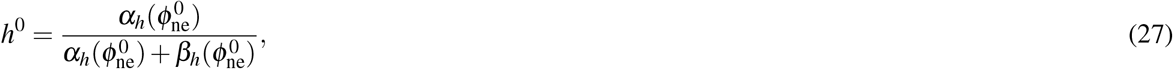

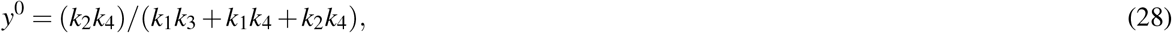

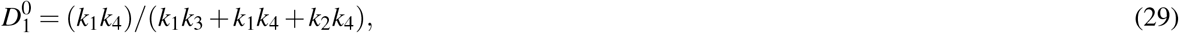

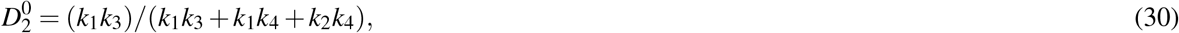

where 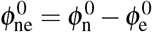.

In the K_ir_4.1 knockout model, a new, steady, initial state is defined by setting the permeability of the glial leak currents (*g*^Na,Leak^ = 2.1 ×10^−2^, 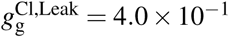), the NaKCl co-transporter (*g*^NaKCl^ = 4.065× 10^−4^), and the glial Na^+^/K^+^-ATPase strength (*I*_g_ = 0.013).

### A.6 Triggering mechanisms

We consider three different mechanisms for triggering CSD:

- Excitatory fluxes: we add excitatory fluxes to the membrane model (24) for the neuron for Na^+^, K^+^ and Cl^−^(i.e. *k* = Na^+^, K^+^, Cl^−^) in the following manner:

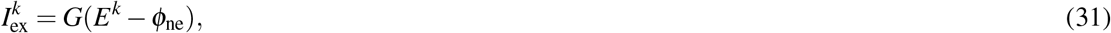

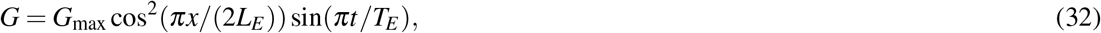

for *t* < *t*_*E*_ and *L*_*E*_ − 2 < *x* < *L*_*E*_, otherwise *G* = 0. We set *L*_*E*_ = 2.0 × 10^−5^ m, *t*_*E*_ = 2 s, and *G*_max_ = 5.0 S/m^2^.
- Topical application of K^+^: we increase the extracellular K^+^ and Cl^−^ in the first 0.001 m (1mm) of the computational domain. In particular, we set the initial concentrations of K^+^ and Cl^−^ such that:

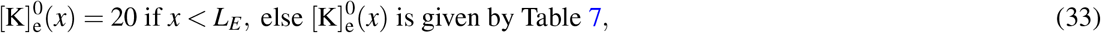

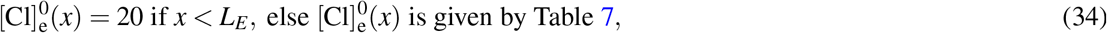

with *L*_*E*_ = 0.001 m.
- Disabled Na^+^/K^+^-ATPase: we disable the neuron and glial pumps in the first 1 mm of our domain.

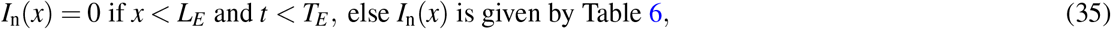

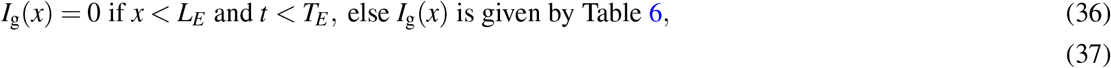

with *L*_*E*_ = 0.001m and *t*_*E*_ = 2s and where 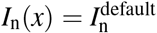 and 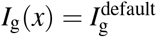 are the default values for the maximum neuronal and glial pump rate listed in, respectively.

## B Supplementary Figures

**Figure 6.**
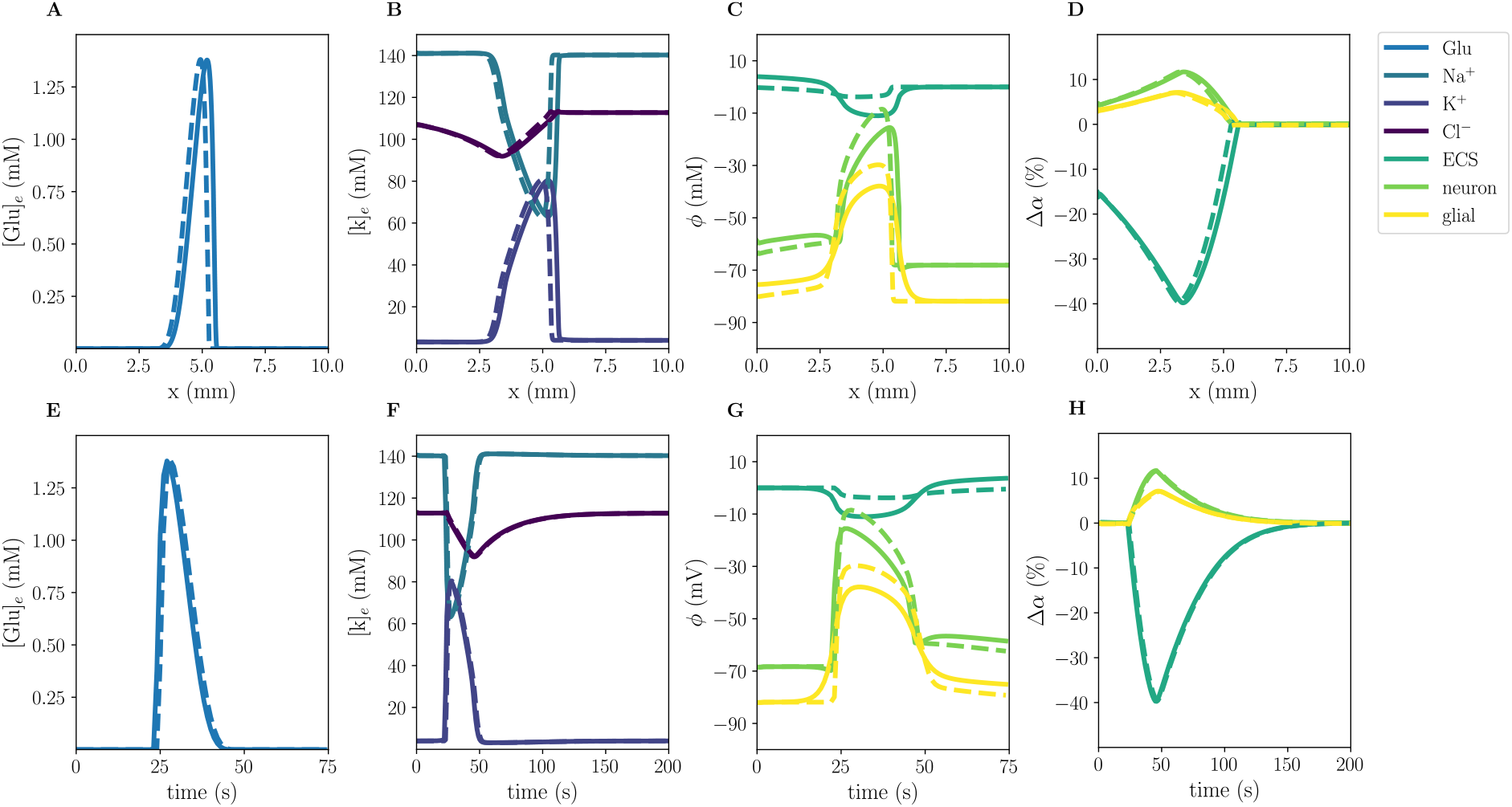
Comparison of model A (solid) and model B (stippled) CSD wave. The upper panels display snapshots of ECS glutamate (**A**), ECS ion concentrations (**B**), potentials (**C**), and change in volume fractions (**D**) at 60 s. The lower panels display time evolution of ECS glutamate (**E**), ECS ion concentrations (**F**), potentials (**G**) and change in volume fractions (**H**) evaluated at *x* = 2.0 mm.

**Figure 7.**
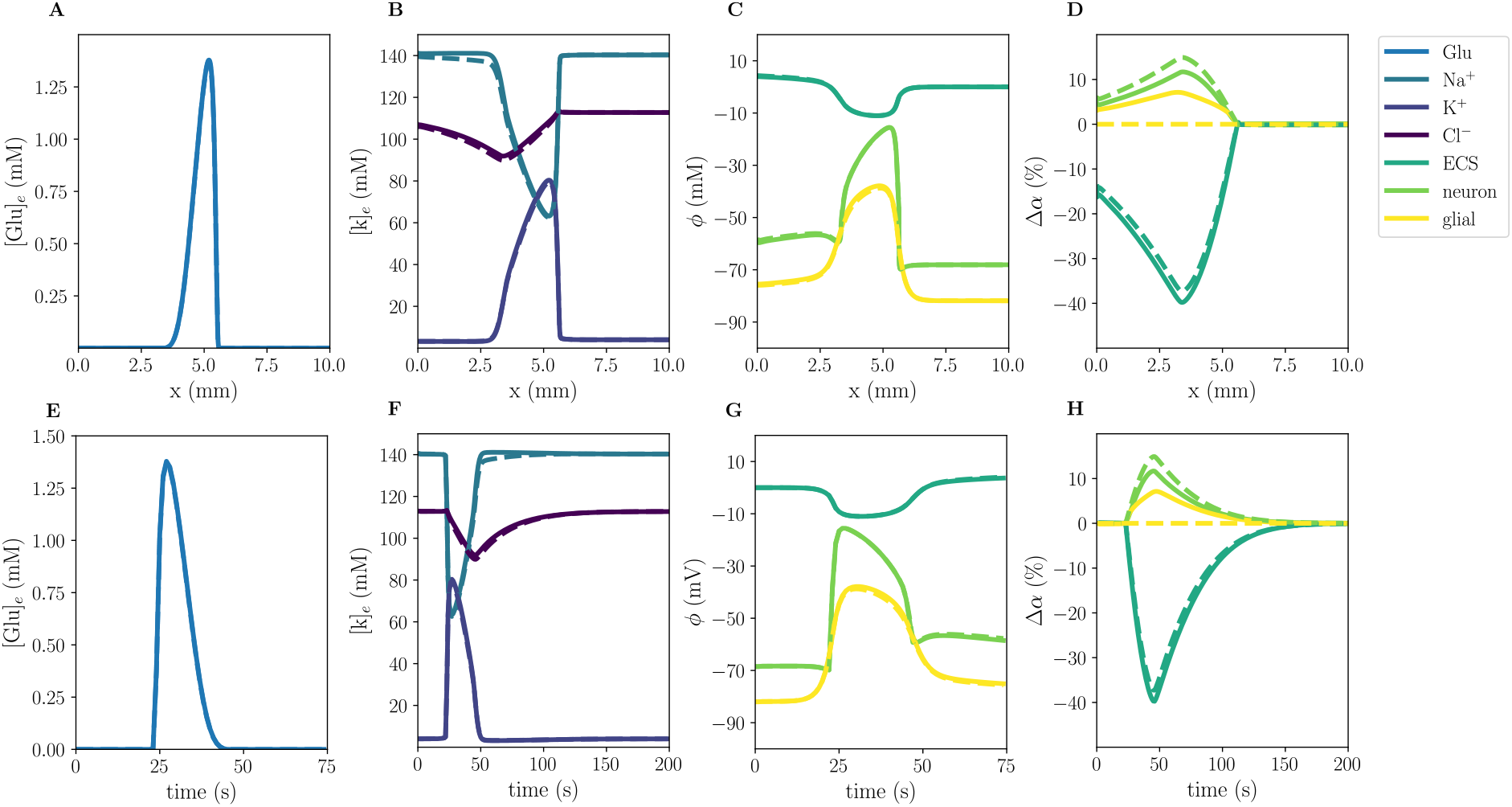
Comparison of model A (solid) and model D (stippled) CSD wave. The upper panels display snapshots of ECS glutamate (**A**), ECS ion concentrations (**B**), potentials (**C**), and change in volume fractions (**D**) at 60 s. The lower panels display time evolution of ECS glutamate (**E**), ECS ion concentrations (**F**), potentials (**G**) and change in volume fractions (**H**) evaluated at *x* = 2.0 mm.

